# Unraveling the *in planta* growth of the plant pathogen *Ralstonia pseudosolanacearum* by mathematical modeling

**DOI:** 10.1101/2024.07.18.604046

**Authors:** Caroline Baroukh, Léo Gerlin, Antoine Escourrou, Stéphane Genin

**Affiliations:** LIPME, Université de Toulouse, INRAE, CNRS, Castanet-Tolosan, France

**Keywords:** Macroscopic modeling, xylem mimicking media, tomato xylem sap, growth dynamics, *R. solanacearum*

## Abstract

*Ralstonia pseudosolanacearum*, a plant pathogen responsible for bacterial wilt in numerous plant species, exhibits paradoxical growth in the host by achieving high bacterial densities in xylem sap, an environment traditionally considered nutrient-poor. This study combined *in vitro* experiments and mathematical modeling to elucidate the growth dynamics of *R. pseudosolanacearum* strain GMI1000 within plants. To simulate the xylem environment, a tomato xylem-mimicking medium containing amino acids and sugars was developed to monitor the growth kinetics of *R. pseudosolanacearum*. Results indicated that glutamine is the primary metabolite driving bacterial growth, while putrescine is abundantly excreted, and acetate is transiently produced and subsequently consumed. A mathematical model was constructed and calibrated using the *in vitro* data. This model was employed to simulate the evolution of bacterial density and xylem sap composition during plant infection. The model accurately reproduced *in planta* experimental observations, including high bacterial densities and the depletion of glutamine and asparagine. Additionally, the model estimated the minimal number of bacteria required to initiate infection, the timing of infection post-inoculation, the bacterial mortality rate within the plant, and the rate at which excreted putrescine is assimilated by the plant. The findings demonstrate that xylem sap is not as nutrient-poor and can sustain high bacterial densities. The study also provides an explanatory framework for the presence of acetate and putrescine in the sap of infected xylem and give clues as to the role of putrescine in the virulence of *R. pseudosolanacearum*.

## Introduction

*Ralstonia pseudosolanacearum* is a gram-negative beta-proteobacteria plant pathogen causing bacterial wilt, a plant disease affecting many important agricultural crops such as tomato, banana, and potato. This bacterium can survive in the soil or water for long periods, enters the host through the roots, begins to multiply in the root apoplasm before migrating to xylem vessels, where an intense multiplication occurs, leading to the plant death through xylem occlusion (1).

Since the 1990s, many studies have focused on the pathogenic determinants of this plant pathogen, such as type-III effectors (2), but very few have examined the metabolic determinants required for nutrient assimilation and efficient multiplication during plant infection (3–8). In particular, it is still not fully understood how *R. pseudosolanacearum* thrives in xylem vessels, an environment usually qualified as nutrient-poor in literature (9), where this pathogen reaches such abundant densities (ten billion cells per gram of plant tissue) that the infected xylem sap appears white (6). Recent studies provided initial insights by highlighting the role of amino acids in xylem sap, particularly glutamine and asparagine (6) (7). Baroukh et al. (7) also demonstrated that xylem sap is not as nutrient-poor as commonly stated if considered as a continuous flow of incoming nutrients for the bacteria. Another unanswered question about *R. pseudosolanacearum* metabolic behavior is the role of the large-scale excretion of putrescine, a polyamine observable both *in vitro* (3, 7) and *in planta* (4, 6). It remains unclear whether this polyamine has a role in virulence (4) and/or bacterial competition (10).

To elucidate *R. pseudosolanacearum* growth in the plant xylem vessels, we developed an *in vitro* xylem-mimicking medium (XMM) and monitored a growth kinetics of *R. pseudosolanacearum* in this medium. In parallel, we developed a mathematical model predicting bacterial growth, substrate consumption, and putrescine excretion by the bacteria. The model was calibrated using experimental data from both *in vitro* growth in XMM and part of the *in planta* kinetics performed by Gerlin et al. (6).This approach was then used to better understand *R. pseudosolanacearum* growth *in planta*.

## Results

### Design of a synthetic xylem-mimicking medium to monitor *Ralstonia* growth

To better assess growth of *Ralstonia pseudosolanacearum* in the xylem sap trophic environment, we followed dynamically GMI1000 strain growth in shake-flasks, on a XMM we created (Table I). Shake-flask experiments allow the sampling of media at different time point and thus the detailed characterization of substrate consumption and metabolites excretion. To create the XMM, we assumed that carbon was the limiting element. The XMM was designed from the classical minimal medium of *R. pseudosolanacearum* containing ions (7), supplemented with trace elements as well as eleven amino acids and two sugars, whose relative composition mimics that of a tomato xylem. To better assess *R. pseudosolanacearum* growth, XMM was more concentrated than tomato xylem sap, with a total carbon content of 50mM. The shake-flask experiments were performed in triplicate, with regular sampling depending on the kinetics. At each sampling time, optical density at 600nm was measured, media was filtrated and its composition was quantified using NMR ^1^H. This approach allowed monitoring of the dynamic behaviors of biomass concentration, substrate concentrations, as well as putrescine concentration, a metabolite excreted abundantly by strain GMI1000 (11).

**Table 1:** Xylem Mimicking Media organic composition and comparison with tomato xylem sap composition (from (6)). Concentrations were measured using NMR in triplicates.

| Molecule | Concentration (mMC) | %age | %age in (6) |
| --- | --- | --- | --- |
| Glutamine | 35.86 ± 1.23 | 71.1%± 2.4% | 74.5% ± 5.9% |
| Lysine | 2.26 ± 0.06 | 4.5% ± 0.1% | 3.3% ± 1.4% |
| Asparagine | 2.15 ± 0.05 | 4.3% ± 0.1% | 3.2% ± 0.8% |
| Arginine | 0.96 ± 0.03 | 1.9% ± 0.1% | 2.6% ± 1.2% |
| Proline | 1.02 ± 0.03 | 2.0% ± 0.1% | 1.5% ± 0.6% |
| Leucine | 1.30 ± 0.04 | 2.6% ± 0.1% | 2.4% ± 0.3% |
| Valine | 0.82 ± 0.02 | 1.6% ± 0.0% | 1.9% ± 0.3% |
| Isoleucine | 0.67 ± 0.02 | 1.3% ± 0.0% | 1.4% ± 0.3% |
| Phenylalanine | 0.69 ± 0.03 | 1.4% ± 0.1% | 1.5% ± 1.0% |
| Threonine | 0.45 ± 0.01 | 0.9%± 0.0% | 1.2% ± 0.3% |
| Tyrosine | 0.38 ± 0.01 | 0.7% ± 0.0% | 1.0% ± 0.4% |
| Glucose | 2.50 ± 0.09 | 4.9% ± 0.2% | 0.4% ± 1.0% |
| Sucrose | 1.00 ± 0.06 | 2.0%± 0.1% | 1.6% ± 1.0% |
| Aspartate | 0.00 ± 0.00 | 0.0% ± 0.0% | 0.3% ± 0.3% |
| Glutamate | 0.00 ± 0.00 | 0.0% ± 0.0% | 0.6% ± 1.3% |
| Alanine | 0.00 ± 0.00 | 0.0% ± 0.0% | 0.2% ± 0.1% |
| Fumarate | 0.00 ± 0.00 | 0.0% ± 0.0% | 0.1% ± 0.2% |
| Ethanol | 0.00 ± 0.00 | 0.0% ± 0.0% | 2.3% ± 4.0% |

GMI1000 growth kinetics observed were consistent with classical bacterial growth curves, comprising a lag phase, an exponential phase and a stationary phase (Fig. 1, Fig S1). Growth stopped at 15 hours, corresponding to the exhaustion of glutamine, the majorly present substrate of the media. At this time point, nearly all the others components were exhausted, except proline, lysine, asparagine and arginine, for which few quantities remained (Fig 1, Fig S1). These results are in agreement with what was already obtained by Baroukh et al. (7) who showed that all metabolites present in XMM could be assimilated either as carbon or nitrogen source. Despite the presence of numerous substrates, no diauxic shift or catabolic repression were observed: GMI1000 catabolized all substrates present in the media at the same time. These results underline that GMI1000 growth is well-adapted to tomato xylem sap, since all tested metabolites were assimilated and few metabolites were remaining at the end of the growth. Regarding excreted metabolites, we observed excretion of putrescine in large quantities, as experimentally observed several times either *in vitro* or *in planta* (3, 4, 6, 7, 12). Interestingly, we also observed the excretion of acetate and its subsequent assimilation.

**Figure 1:**
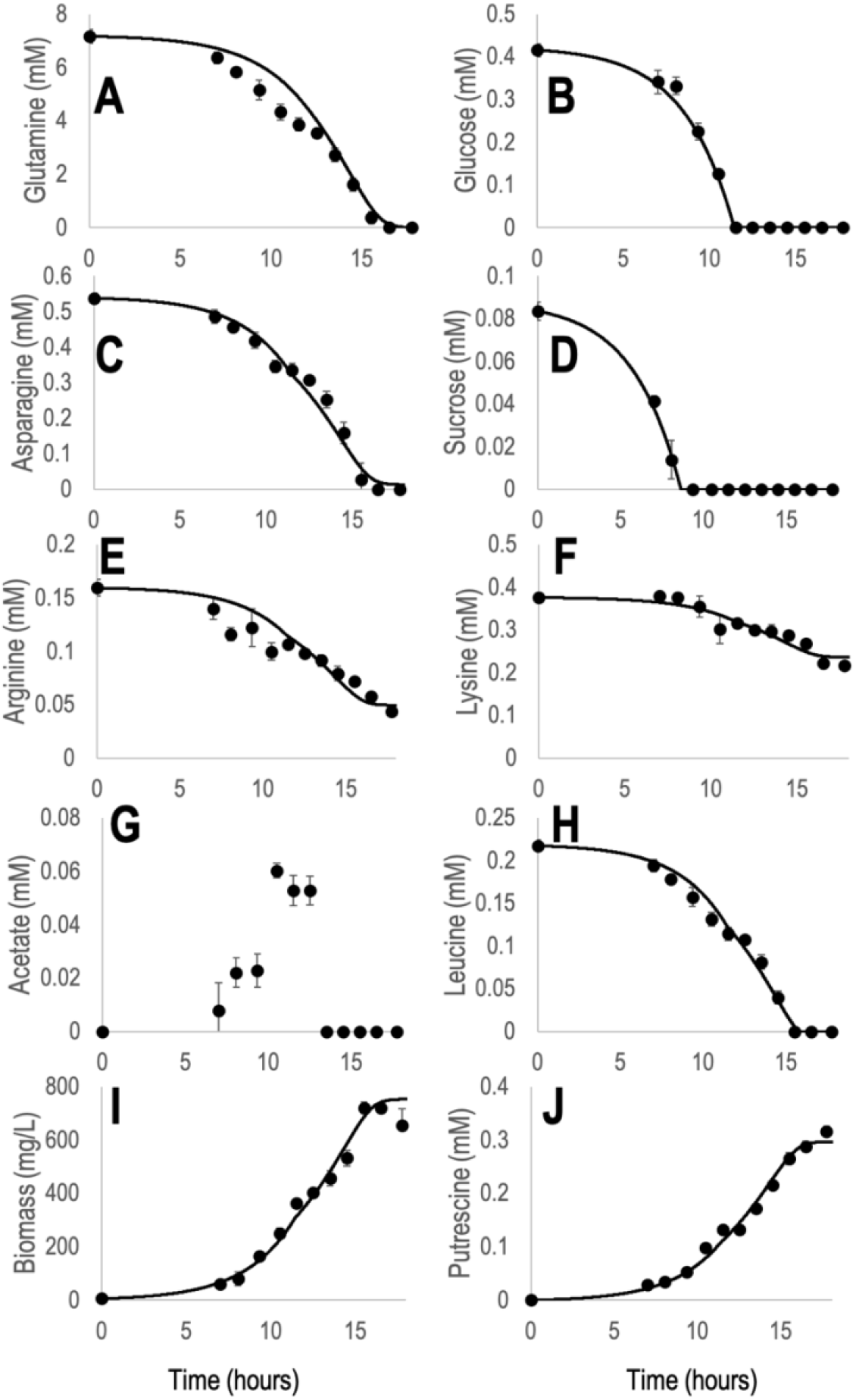
Growth monitoring of *R. pseudosolanacearum* GMI1000 strain in a xylem mimicking media. Dots are experimental data, line represents model’s simulation results. Metabolites concentration over time was assessed by NMR analysis of culture filtrate. Biomass concentration over time was assessed by optical density at 600nm. A. Glutamine. B. Glucose. C. Asparagine. D. Sucrose. E. Arginine. F. Lysine. G. Acetate. H. Leucine. I. Biomass J. Putrescine. Other metabolites dynamics are available Fig S1.

### Modeling Ralstonia growth in vitro

To better understand *R. pseudosolanacearum* growth in xylem mimicking media and estimate several important parameters such as its growth rate, we built a mathematical model using a parsimonious number of parameters implying a Monod kinetic for glucose and glutamine substrates, and biomass proportional assimilation rates for all other substrates. Putrescine excretion was also assumed proportional to biomass growth. As we do not model *R. pseudosolanacearum* cell metabolism but only its macroscopic behavior, we did not predict acetate excretion and assimilation (see discussion section). From these kinetic rates, a mass balance was performed to derive ordinary differential equations predicting substrates, putrescine and biomass dynamics along time. The choices regarding the mathematical equations were made by observing the consumption kinetics of all substrates: glutamine, the most abundant metabolite and presumably the main carbon substrate driving growth, as the assimilation of other substrates stops upon glutamine exhaustion, with the exception of glucose, which maintains its own assimilation rate.

The total number of parameters for the model thus comprised 19 parameters, to represent the dynamic behavior of 13 substrates, putrescine and biomass over time (Table 2). Parameters estimations were then performed using experimental data for shake-flask experiments and an optimization algorithm minimizing the squared error between experimental data and model’s simulation. Overall, the calibrated mathematical model could predict correctly the consumption of the 13 substrates as well as biomass production and putrescine excretion (Fig 1, Fig S1). The parameters allowed to have an estimation of *R. pseudosolanacearum* growth rate in xylem mimicking media as well as substrate assimilation rate and putrescine excretion rate.

**Table 2:** Models parameters. Parameters estimations was performed using a L-BFGS algorithm using scipy library in python 3.7. All parameters were estimated for XMM except for the mortality rate and putrescine uptake rate which was estimated using experimental data of Gerlin et al (6)

| Parameter (unit) | Value | Parameter (unit) | Value |
| --- | --- | --- | --- |
| $k_{gln}$ (mM.gB<sup>-1</sup>) | $1.11 * 10^{-2}$ | $k_{arg}$ (mM.gB<sup>-1</sup>) | $1.47 * 10^{-4}$ |
| $k_{putr_{gln}}$ (mM.gB<sup>-1</sup>) | $4.21 * 10^{-4}$ | $k_{sucr}$ (mM.gB<sup>-1</sup>) | $7.43 * 10^{-4}$ |
| $\mu_{max_{gln}}$ (h<sup>-1</sup>) | 0.26 | $k_{tyr}$ (mM.gB<sup>-1</sup>) | $1.39 * 10^{-4}$ |
| $K_{S_{gln}}$ (mM) | 0.85 | $k_{pro}$ (mM.gB<sup>-1</sup>) | $9.61 * 10^{-4}$ |
| $k_{gluc}$ (mM.gB<sup>-1</sup>) | $4.13 * 10^{-3}$ | $k_{lys}$ (mM.gB<sup>-1</sup>) | $1.87 * 10^{-4}$ |
| $k_{putr_{gluc}}$ (mM.gB<sup>-1</sup>) | $2.42 * 10^{-4}$ | $k_{thr}$ (mM.gB<sup>-1</sup>) | $1.13 * 10^{-4}$ |
| $\mu_{max_{gluc}}$ (h<sup>-1</sup>) | 0.11 | $k_{iso}$ (mM.gB<sup>-1</sup>) | $1.58 * 10^{-4}$ |
| $K_{S_{gluc}}$ (mM) | $1.21 * 10^{-4}$ | $k_{asn}$ (mM.gB<sup>-1</sup>) | $7.02 * 10^{-4}$ |
| $k_{phe}$ (mM.gB<sup>-1</sup>) | $2.86 * 10^{-4}$ | $k_{arg}$ (mM.gB<sup>-1</sup>) | $1.47 * 10^{-4}$ |
| $k_{val}$ (mM.gB<sup>-1</sup>) | $2.05 * 10^{-4}$ | mortality | $8.52 * 10^{-3}$ |
| $k_{leu}$ (mM.gB<sup>-1</sup>) | $3.11 * 10^{-4}$ | $k_{Upt_{putr}}$ | 0.173 |

### Modeling Ralstonia growth in planta

To simulate the behavior of *R. pseudosolanacearum in planta*, we used the mathematical model calibrated on *in vitro* kinetics, but changed the environmental conditions to those presumably encountered in plants. The model was thus modified using the following assumptions: since xylem sap is a continuous flow of metabolites linked to the plant’s transpiration rate, xylems vessels were considered as a chemostat, where substrates are not only available initially but also continuously supplied. Interpolated transpiration data (6) were used to estimate this flow throughout the model simulation (Fig S2). As experimentally verified by Gerlin et al. (6, 13), it was assumed that in a non-infected plant, xylem sap concentration did not vary significantly over the 8-day infection window. This assumption implied that the incoming metabolite concentration is equal to the concentration measured in a non-infected plant. In addition, plant defense responses cause mortality among the bacteria. This was mathematically translated into a mortality rate proportional to the biomass density as well as the transpiration rate. A linear dependence on the transpiration rate was assumed because it allowed the model to better fit the observed experimental data. From a biological standpoint, this hypothesis implies that there is less mortality when there is less transpiration, which can be explained by the fact that when transpiration decreases, the plant is dying and thus its defenses have been overcome by the pathogen, resulting in less bacterial mortality. To be able to predict correctly putrescine content in infected xylem sap, an uptake term by the plant aerial parts, proportional to the transpiration rate, was also added. Finally, to compare biomass in mg/L, as measured in shake-flask experiments and predicted by the model, and biomass in CFU/gFW, as measured *in planta*, we calculated a conversion factor based on both xylem vessel volume, plant density and plant fresh weight (see Material and Methods section for further details).

The mortality rate and the putrescine uptake by the plant was calibrated using the biomass, glucose, glutamine, asparagine and putrescine kinetic data of infected xylem sap previously determined (6) from day 3 to 7 post infection (Table 2). Besides these two new parameters related to the plant environment, all the others parameters of the models were those calibrated on the *in vitro* kinetic data (Table 2).

### Predicting the dynamic evolution of organic metabolites in infected xylem sap

The model simulations showed that it could correctly predict the dynamic evolution of the organic composition of infected xylem sap along with *R. pseudosolanacearum* density (Fig 2, Fig 3). In particular, glutamine, asparagine, glucose and lysine depletion dynamics were well-predicted (Fig 2, Fig 3). Phenylalanine, valine, sucrose, leucine, isoleucine and tyrosine were depleted faster in the model compared to experimental data, even though model predictions were still within the range of the standard-deviation of these metabolites’ concentrations (Fig 3). For proline, threonine and arginine, the model predicted depletion whereas experimental data at 7 days post infection do not show depletion of these metabolites (Fig 3).

**Figure 2:**
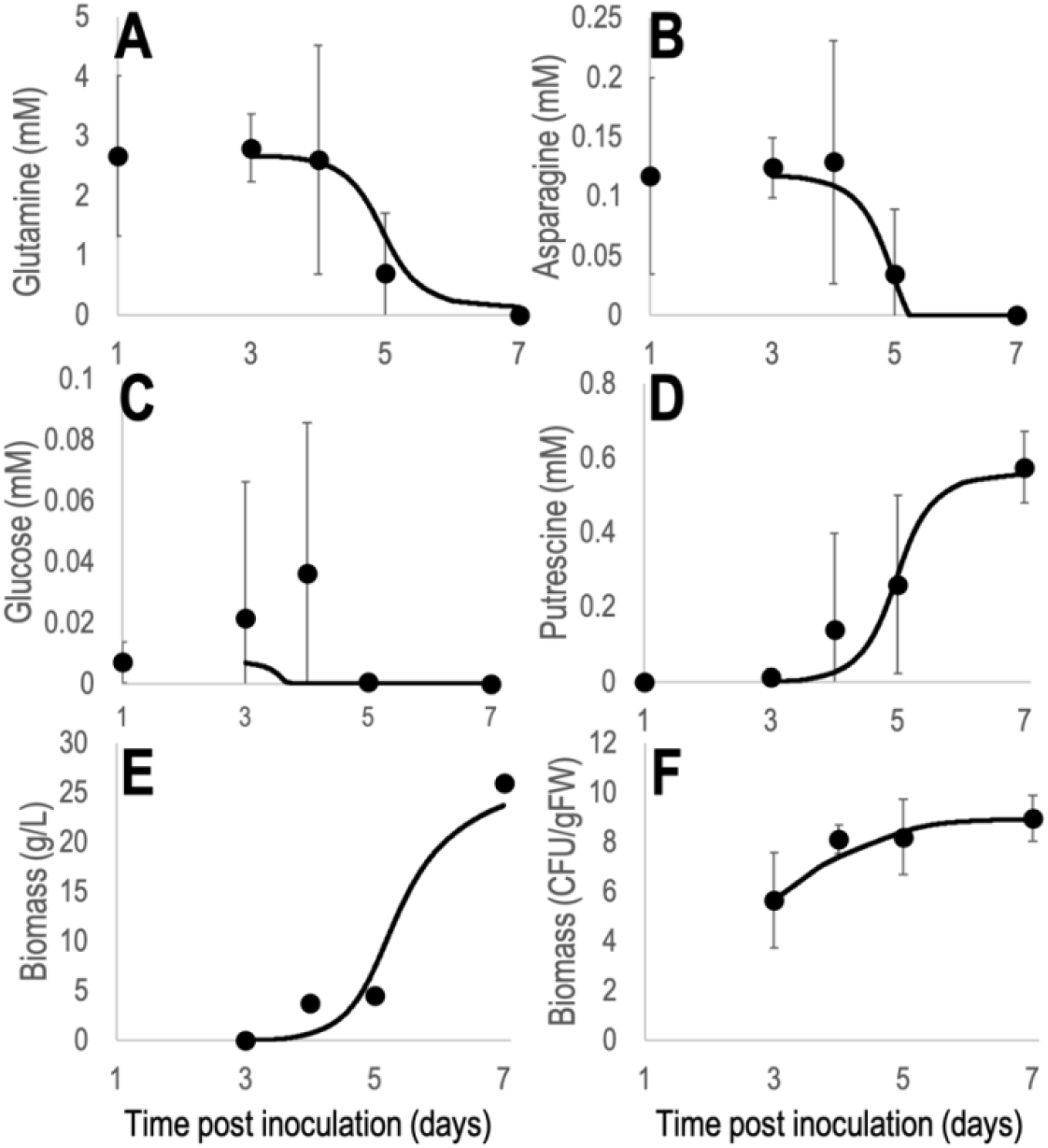
Comparison of glutamine (A), asparagine (B), glucose (C), putrescine (D) and GMI1000 biomass (E and F) *in planta* dynamic behavior between model’s predictions and experimental data. Dots are experimental data taken from Gerlin et al. (6), lines represent model’s predictions.

**Figure 3:**
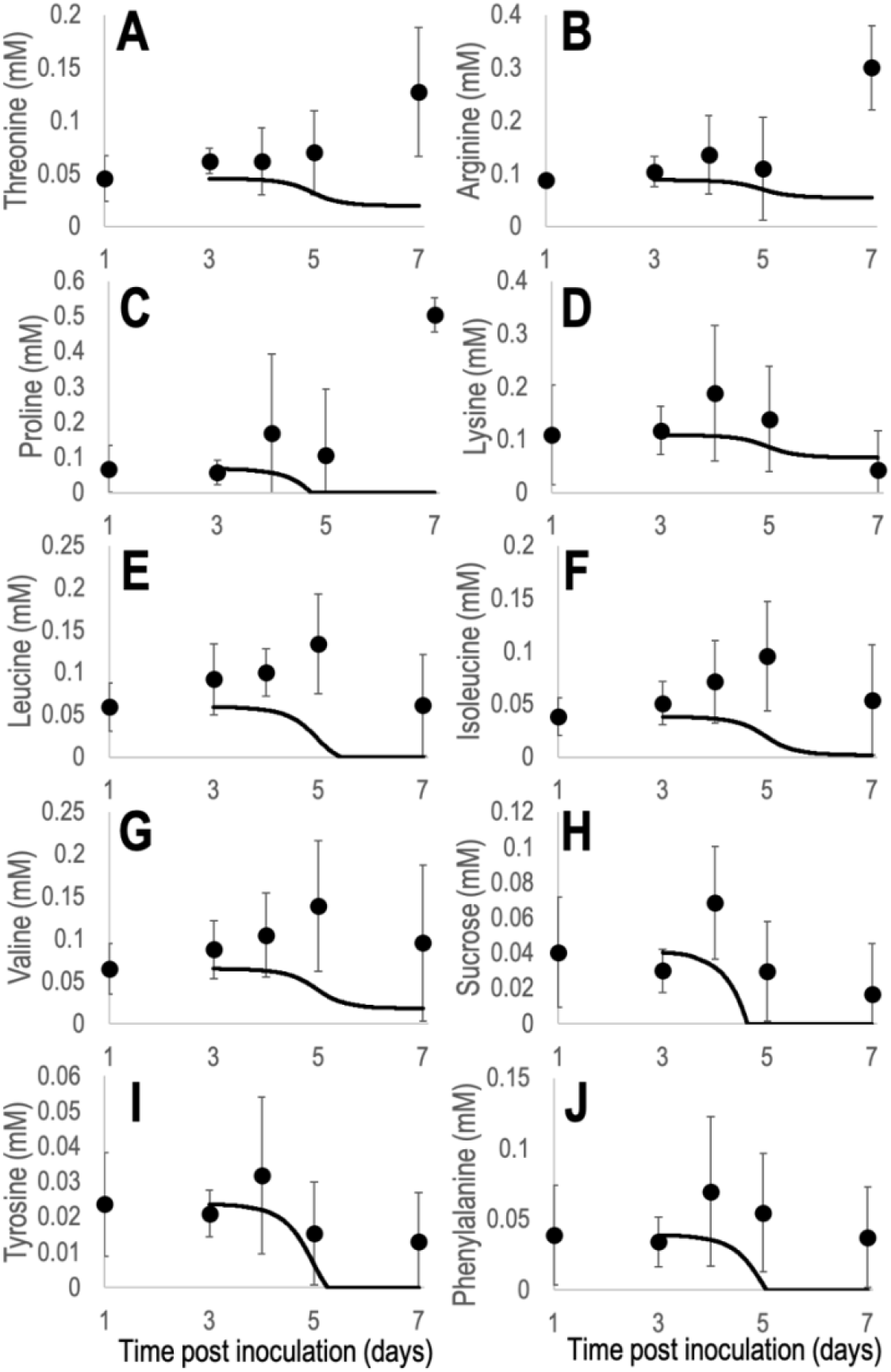
Comparison of xylem metabolites *in planta* dynamic behavior between model’s predictions and experimental data. Dots are experimental data taken from Gerlin et al. (6), lines represent model’s predictions. A. Threonine B. Arginine. C. Proline. D. Lysine. E. Leucine. F. Isoleucine. G. Valine. H. Sucrose. I. Tyrosine. 236 J. Phenylalanine.

### Predicting maximal bacterial densities achievable during infection

In the model, the maximal density achievable by the pathogen is the resulting balance between the mortality rate and xylemic substrate concentrations available for *R. solanacearum* growth rate (mainly glutamine). Substrate concentrations depend themselves on the incoming flux of substrates which is governed by the transpiration rate of the plant. Upon infection, transpiration rate of the plant severely decreases due to xylem occlusion by the bacteria (1) (Fig S1). The model predicted that the maximal biomass density that could be reached was 10^8.92^ CFU/gFW (equiv. 23.67 g/L), close to the value measured experimentally (6). Without a mortality rate, the maximal density that could be reached was larger (10^9.64^ CFU/gFW, equiv 125g/L). In this scenario, the slow-down of *R. solanacearum* growth was only due to i) the exhaustion of metabolites driving its growth and ii) the decrease of incoming metabolites due to the decrease of the transpiration rate of infected plants. If transpiration is set to be constant at 100mL/day, that is the average transpiration observed 3 days post infection for infected and healthy plants, *R. pseudosolanacearum* can achieve a maximal density of 10^9.93^ CFU/gFW (equiv. 247g/L) without any mortality. It thus seems possible that *R. pseudosolanacearum* can reach densities up to 10^10^ CFU/gFW in the plant by growing only on available organic metabolites of xylem sap.

### Predicting putrescine excretion by the bacteria and implications for the host

The mathematical model also predicted putrescine excretion by the bacteria. If putrescine excreted is not uptaken by the plant, putrescine final concentration reaches 32.29mM, 58 times more than what is observed experimentally (Fig 2). Using the model, a putrescine plant uptake term was thus estimated so that the dynamic of putrescine concentration over time is similar to what was observed experimentally. This total uptake of putrescine over time by the plant is far from being negligible: 31.73mM over 4 days.

### Estimating the time required for *R. solanacearum* to reach xylem vessels

To date, it is still unclear how many *R. pseudosolanacearum* founder bacteria cause infection in the plant and how long it takes them to reach the xylem vessels where fast multiplication occurs. The mathematical models developed in this study, which are based on ordinary differential equations, provided an initial answer to these questions. Indeed, to solve the model, initial conditions are necessary. Since biomass concentration was known only from day 3 post inoculation (6), the model was thus simulated and calibrated from day 3. However, once the model calibrated, it can be used to determine the initial number of bacteria infecting the xylem (CFUs) if the start time of the xylem infection is known and vice-versa.

By simulation, we thus determined at which time post inoculation the xylem infection started for different numbers of bacteria initiating the plant infection (Table 3). The model shows that a minimum of 21 bacteria are necessary to reach a biomass of 10^5.66^ CFU/gFW at day 3, as previously determined (6). As expected, the time to infection onset increased with the number of bacteria initiating infection: almost 7.5 hours are necessary before infection onset if 100 bacteria initiate it, up to 20 hours for 1000 bacteria. Since xylem infection is unlikely to start immediately, even though the inoculation was performed on scarified roots (6), the infection onset time is most likely at least 7.5 hours. It should be noted that the prediction of xylem infection start-up time depends on the model’s mortality rate. If there is no bacterial mortality, the bacteria take much longer to infect the plant: if a single bacterium causes the xylem infection, the model predicts a xylem infection start-up time of 22.8 hours, rising to 44 hours for 1000 bacteria.

**Table 3:** Estimation, by model simulation, of the xylem infection start time for different number of bacteria initiating the xylem infection with and without bacterial mortality.

| Number of founders bacteria infecting the xylem (CFU) | Xylem infection start time (hours post inoculation) | Xylem infection start time without mortality (hours post inoculation) |
| --- | --- | --- |
| 1 | Not possible | 22.8 |
| 10 | Not possible | 29.88 |
| 21 | 0 | 32.09 |
| 100 | 7.44 | 36.84 |
| 500 | 16.06 | 41.76 |
| 1000 | 19.92 | 43.87 |

## Discussion

### Xylem mimicking media: an *in vitro* media close to the plant environment

To better assess *R. pseudosolanacearum* growth *in planta*, we designed a xylem mimicking media (XMM) that mimics the organic composition of concentrated plant xylem sap. This medium offers the advantage of simulating the plant carbon trophic environment while providing the benefits of *in vitro* experiments. The concentrated organic carbon content allows for improved monitoring of bacterial growth (OD, metabolite concentrations) compared to ex-vivo xylem sap, which is a nutrient-poor environment supporting limited bacterial multiplications and underestimating the access to nutrients occurring in planta. In addition, XMM facilitates easier sampling of large volumes of filtrate or bacterial biomass compared to ex-vivo sap or *in planta* experiments. Harvesting sufficient quantities of sap or biomass from plants is often challenging, particularly when studying *R. pseudosolanacearum* at biomass densities below the Phc quorum sensing threshold (14). Finally, our designed XMM provides a reproducible trophic environment, which is crucial for comparing experiments performed under various conditions, such as growth of mutant versus wild-type strains or growth of multiple strains. This reproducibility is not achievable with ex-vivo xylem sap, whose composition can vary by up to 30% between individual plants (6, 7, 13).

Currently, the media used to study *R. pseudosolanacearum in vitro* behavior is either a complete medium composed of bactopeptones, casamino acid and yeast extract or a minimal medium composed of essential ions for growth supplemented with glucose or glutamate (15) or more recently glutamine (13). The complete media was already shown to inadequately represent the plant environment, as it fails to induce expression of the type III secretion system (16, 17). A minimal media supplemented with glutamine is considered a good approximation, given that glutamine can constitute up to 73% of the total organic carbon present in xylem sap. However, XMM, which incorporates all main organic compounds found in xylem sap in proportions similar to those in the plant, provides an even more accurate representation of the *in planta* trophic environment. Our mathematical model, calibrated using data from this medium, further supports the validity of XMM. Indeed, the model’s ability to correctly predict the depletion kinetics of most metabolites in infected plants proves that XMM closely mimics the trophic environment encountered by the bacteria in the plant.

### Role of excreted metabolites by *R. pseudosolanacearum*

When growing on XMM, we showed that *R. pseudosolanacearum* excreted large quantities of putrescine as well as acetate which was later on re-assimilated. Interestingly, infected tomato xylem sap is enriched in both those molecules (6).

### Putrescine

Excretion of putrescine in XMM is in line with previous experimental results which showed that this polyamine is essential for *R. pseudosolanacearum* growth and is excreted by the bacteria into large quantities in synthetic media (3, 4, 7) and during plant infection (5). Putrescine seems to be excreted in a constitutive manner, being produced in all the *R. pseudosolanacearum* growth media tested to date (7). The role of this excreted putrescine for *R. solanacearum* is unclear. On one hand putrescine pre-treated plants had increased sensibility to the pathogen (4), and on the other hand mutant defective for *phcA*, a master regulator of virulence functions, excretes putrescine in larger quantities than the wild-type strain (3). These observations raise the question of whether putrescine has a role in virulence or is merely a by-product of growth. In this study, the mathematical model showed that putrescine needs to be up-taken in large quantities by the plant so that models’ predicted concentration in xylem sap fits with published experimental data (6). The role of putrescine produced by bacteria could be to disrupt polyamine metabolism in the plant. An external source of putrescine might lead to the inhibition of putrescine-related synthesis reactions and catalysis of putrescine-related degradation reactions. For example, the metabolic reaction converting spermidine to putrescine could be inhibited, thus decreasing the plant’s H_2_O_2_ production, which is important in the ROS response (12). Given the design of our mathematical model, which represents the plant only as a chemostat and not as an organism in itself, it is difficult to conclude whether the uptaken quantity is large enough to disrupt plant polyamine metabolism in this way. However, a metabolic model incorporating both the plant and the pathogen should help to answer such question.

### Acetate

Acetate is a product of cell fermentation, and its excretion is observed, for example, when *E. coli* grows rapidly in culture media, in particular if several carbon sources are available (18). This behavior is known as overflow metabolism and is often described as an inefficient use of metabolic routes and apparent waste of energy resources. Its origin lies in the presence of limiting metabolic pathways in the cell, such as the TCA cycle or the respiratory chain compared to the assimilation and catabolism of substrates. Given the relatively rapid growth of *R. pseudosolanacearum* GMI1000 in XMM (growth rate around 0.30h^-1^ when both glucose and glutamine are present) and the high cell carbon input in this case (simultaneous assimilation of 13 compounds), acetate excretion and subsequent assimilation are likely due to a similar phenomenon of metabolic overflow. Interestingly, Gerlin et al. (6) observed the appearance and subsequent decrease of acetate in the xylem sap of in infected tomato plants, which aligns with what is observed *in vitro* in XMM.

### 3-hydroxybutyrate

The appearance and subsequent decrease of 3-hydroxybutyrate (3HB) in infected tomato xylem sap was also reported (6). 3HB is the monomer of polyhydroxybutyrate (PHB), a carbon storage molecule largely accumulated in *R. solanacearum* (19, 20). However, the appearance of this metabolite has not been observed in XMM. Thus, its production appears to be linked to *in planta* conditions or a stress not present in our shake-flask experiments. Interestingly, *R. pseudosolanacearum* has been shown to assimilate 3HB to sustain its growth (3, 8). Therefore, the observed decrease in 3HB in infected xylem sap is likely due to *R. pseudosolanacearum* consumption. However, the question arises whether *R. pseudosolanacearum* or the plant is at the origin of this 3HB production. Notably, 3HB can be produced both by tomato plants and *R. pseudosolanacearum* (3, 13). In particular, *R. pseudosolanacearum* possesses an extracellular PHB depolymerase (3), which could depolymerize PHB present in infected xylem sap. The source of this PHB could be the lysis of dead *R. pseudosolanacearum* bacteria. To evaluate the plausibility of this hypothesis, the mortality rate from the mathematical model was used to assess the potential quantity of PHB in infected xylem sap. Assuming a 23% PHB content in bacterial biomass, as measured by Alves et al (20), the quantity of PHB as 3HB equivalent in xylem sap could reach 105mM. This is approximately 297 times higher than what was observed experimentally by Gerlin et al (6). Thus, even if only a tiny fraction of PHB from bacterial lysis is depolymerized and 3HB is assimilated by *R. pseudosolanacearum*, it would yield quantities of 3HB comparable to those detected in infected xylem sap. Therefore, it is plausible that the observed 3HB could originate from the depolymerization of PHB resulting from bacterial lysis in infected xylem sap.

### Xylem sap is sufficient to sustain high bacterial densities

Usually, the maximal density reached by infecting *R. solanacearum* populations in the plant is around 10^9^ CFU/gFW -10^10^ CFU/gFW (21, 22), which gives an equivalent dry weight of 28.57 -285.7g.L^-1^. Xylem sap is often described in literature as a nutrient-poor environment (9), which seems paradoxical given the high bacterial densities it can support. However, xylem sap composition is typically measured at a single time point, not as a continuous flow of metabolites. As already discussed by Baroukh et al. (7), when considering xylem as a flow of metabolites, xylem is far from nutrient-poor and can deliver up to 8M of carbon over three days. In this study, the mathematical model demonstrated that even with a bacterial mortality and decreasing plant transpiration rates, *R. pseudosolanacearum* can reach bacterial densities to up to 10^9^ CFU/gFW by growing solely on substrates available in xylem sap, without any modification of plant metabolism or necrotrophic behavior. This high density is limited by both substrate exhaustion by *R. pseudosolanacearum* and drastically reduced access to substrates due to decreased transpiration. In this scenario, *R. pseudosolanacearum* therefore limits its own growth by clotting xylem vessels, thereby impeding the xylem flow that would otherwise supply it with nutrients.

When the bacterial mortality and the decrease in transpiration rate are removed from the model, it predicts a final bacterial density close to 10^10^ CFU/gFW. In this case, *R. pseudosolanacearum*’s growth is limited solely by substrate exhaustion: its population becomes too high to sustain itself on the available xylem flow. This density of 10^10^ CFU/gFW thus appears to be the maximal achievable density given the xylem composition and flow in the tomato plants (6). To achieve higher densities, the xylem composition would need to be more concentrated. For instance, the mathematical model indicates that the xylem would need to be 20 times more concentrated for *R. pseudosolanacearum* to reach a bacterial density of 10^11^ CFU/gFW.

## Conclusion

In this study, the use of XMM, combined with mathematical modeling, has helped elucidate *R. pseudosolanacearum in planta* growth, particularly explaining how such high bacterial densities can be achieved in a seemingly nutrient-poor environment. XMM also provided insights into the presence of both acetate and putrescine in infected xylem sap, while the absence of 3HB suggests it may be a plant condition-specific metabolite (6). In the future, XMM could be a valuable tool for various *in vitro* experiments to better understand *R. pseudosolanacearum* biology. These could include transcriptomics, proteomics, metabolomics or microfluidic studies (23), especially in cases where *in planta* experiments are challenging to conduct. On the modeling side, a metabolic model representing metabolism of both the plant and the pathogen would allow to better understand the metabolic interactions happening during a xylem infection including, among others, if putrescine assimilation by the aerial part of the plant is enough to disrupt polyamine metabolism in the plant.

## Material and Methods

### Bacterial strains and inoculum

The *R. pseudosolanacearum* strain used for this study is the wild-type strain GMI1000 (24). The bacteria were plated on complete medium (with g.L^−1^: Bactopeptone: 10; casamino acid: 1; yeast extract: 1), supplied with glucose 20% (5 g.L^−1^) and TTC (0.05 g.L^−1^) for 48 h at 28°C. Then, a bacterial colony was grown in liquid complete medium. Inocula were shaken at 180 rpm on an orbital shaker placed in an incubator at 28°C.

### Shake-flasks growth experiments

GMI1000 was grown in 1L shake flasks shaken at 180 rpm on an orbital shaker placed in an incubator at 28°C. The starting volume culture was 250mL. Culture medium was a xylem mimicking media, composed of a minimal medium (with g.L^−1^: FeSO_4_,7H_2_O: 1.25 × 10^−4^, (NH4)_2_,SO_4_: 0.5, MgSO_4_,7H_2_O: 0.05; KH_2_PO_4_: 3.4) supplemented by trace elements (with mg.L^−1^: FeSO_4_,7H_2_O: 5, Na_2_EDTA,2∙H_2_O: 50, ZnSO_4_,7∙H_2_O: 22, H_3_BO_3_: 11.4, MnCl_2_,4∙H_2_O: 5.04, CoCl_2_,6∙H_2_O: 1.6, CuSO_4_,5∙H_2_O: 1.56, (NH_4_)_6_Mo_7_O_24_,4∙H_2_O: 1.12) and amino acids and sugars as carbon sources for a total of 50mMC (see Table I for exact composition). pH was adjusted to 6.5 using KOH.

Dynamic sampling was performed to monitor biomass growth (OD measurement at 600nm using a spectrometer), substrate consumption (NMR analysis) and products excretion (NMR analysis). Sampling time was chosen according to the kinetic evolution with intensive sampling during the exponential growth phase. After OD measurement, the collected volume for each time point was filtered with a 0.22 µm micro-filter and conserved at -20°C for further analysis.

### NMR analysis of culture filtrate

Small-size organic metabolites from the liquid culture in shake flaks were quantified by NMR. Samples were analyzed by 1D ^1^H NMR on MetaToul analytics platform (UMR5504, UMR792, CNRS, INRAE, INSA 135 Avenue de Rangueil 31077 Toulouse Cedex 04, France), using the Bruker Advance 500 MHz. The samples were kept all along the analysis at of 280°K. TSP-d4 standard (Sodium 3-(trimethylsilyl)(1-13C,2H4)propanoate) was used as a reference. Resonances of metabolites were manually integrated using Bruker TopSpin software and the concentrations were calculated based on the number of equivalent protons for each integrated signal and on the TSP final concentration.

### Construction of a mathematical model predicting *R. pseudosolanacearum* growth in synthetic xylem media

The mathematical model predicting GMI1000 growth on xylem mimicking media was built using a mass-balance and by assuming Monod kinetic rates on glutamine and glucose. For the others substrates, assimilation was assumed to be proportional to biomass. This yielded the following mathematical equations:

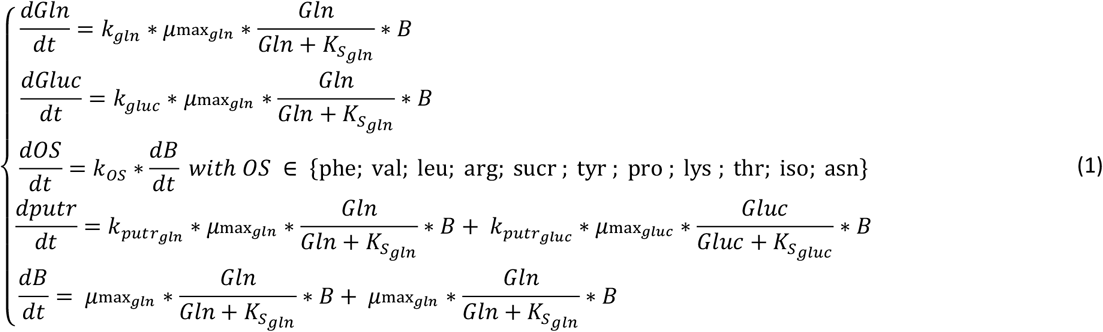

With:

- *Gln, Gluc, putr* and *OS* the concentrations (mM) of respectively glutamine, glucose, putrescine and the other substrates (phenylalanine, valine, leucine, arginine, sucrose, tyrosine, proline, lysine, threonine, isoleucine, asparagine)
- *B*, the *R. pseudosolanacearum* GMI1000 biomass concentration (in mg.L^-1^)
- *kgln* (mM.gB^-1^) the quantity of glutamine necessary to yield 1g of biomass, *μ*max_*gln*_ (h^-1^) the maximal growth rate on glutamine, 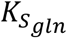 (mM) the semi-saturation growth term on glutamine, 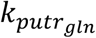 (mM.gB^-1^) the quantity of putrescine excreted per gram of biomass when growing on glutamine
- *kgluc* (mM.gB-1) the quantity of glucose necessary to yield 1g of biomass, *μ*max_*gluc*_ (h^-1^) the maximal growth rate on glucose, 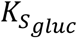 (mM) the semi-saturation growth term on glucose, 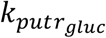 (mM.gB^-1^) the quantity of putrescine excreted per gram of biomass when growing on glucose
- *kOS* (mM.gB^-1^) the quantity of other substrates (phenylalanine, valine, leucine, arginine, sucrose, tyrosine, proline, lysine, threonine, isoleucine, asparagine) assimilated by unit of biomass when growing on a xylem synthetic media.

Model parameters were estimated by minimizing the squared error between experimental data (biomass and composition of the media over time) and models predictions using a L-BFGS-B algorithm (scipy.optimize python library). Optimization was launched multiple times with different initial conditions to reduce the risk of finding local minima. The parameter set fitting the best experimental data was chosen.

### Adaptation of the mathematical model to *in planta* conditions

To adapt the model of *R. solanacearum* growing on synthetic xylem to *in planta* conditions, addition of a dilution term (*D*, h^-1^) and incoming substrate terms (*Ksi*_*in*_, mM) for all substrates were added to the model. Indeed, xylem sap can be seen as a continuous flow of nutrients whose rate is equivalent to the plant transpiration rate. Thus equations’ system (1) is transformed to:

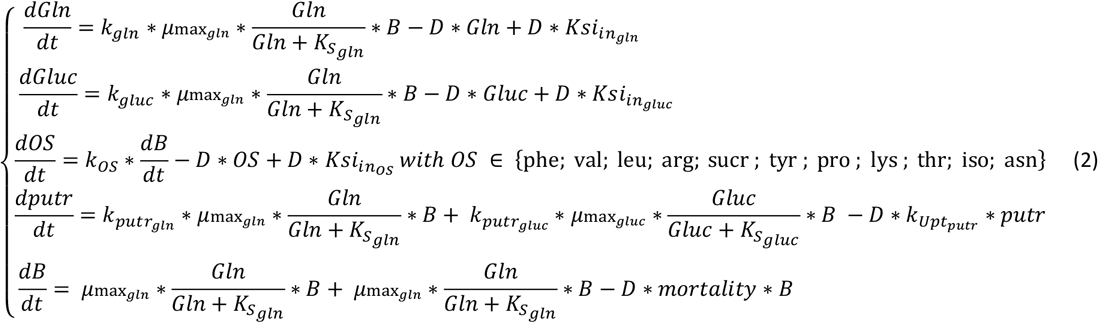

The incoming substrate terms (*Ksi*_*in*_) was assumed equal to the concentration measured in a non-infected plant. Dilution rate D was computed by interpolating the transpiration rate of the plant (6) and divided it by the xylem volume, which was estimated to be 1.063% of the plant volume (7), which itself was estimated using the plant fresh weight (also interpolated from (6)) to the plant density:

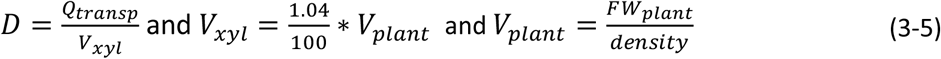

*R. pseudosolanacearum* biomass was assumed not submitted to this dilution rate since *R. pseudosolanacearum* forms biofilm inside the xylem vessels. However, to better predict the experimental data observed *in planta* (6), a *mortality* rate, proportional to the plant transpiration rate (and thus its infection state) was assumed. Finally, to better fit putrescine concentration in infected xylem sap, an uptake rate 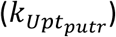, proportional to the plant transpiration rate (and thus its infection state) was also assumed. Mortality and putrescine uptake parameters were estimated by minimizing the squared error between models predictions and experimental data of (6) composed of glutamine, glucose, asparagine, biomass and putrescine dynamic evolution from day 3 to 7.

### Estimation of 3HB accessible by depolymerization of PHB issued from bacterial lysis

According to Alves et al (20), 23 and 45% of *Ralstonia pseudosolanacearum* dry weight is composed of P(3HB). The potential quantity of 3HB to which *Ralstonia pseudosolanacearum* has access is thus:

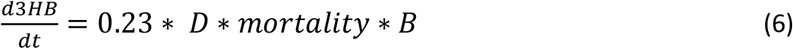

### Conversion of CFU/gFW into OD_600nm_ and mg/L of biomass

Conversion of CFU/gFW into OD_600nm_ and mg/L of biomass is necessary since *R. pseudosolanacearum* biomass was measured in CFU/gFW in (6) and we measured *R. pseudosolanacearum* biomass as OD and equiv. mg/L in xylem mimicking media. As explained in (7), from supplementary Table S4 from Peyraud et al. (3), we have access to conversion between CFU/mL and OD, and OD to mg/L of biomass, using the following formulae:

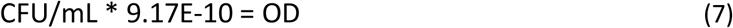

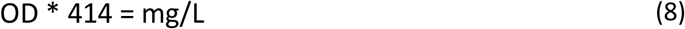

In addition, thanks to the estimation of the volume of the xylem, we can obtain conversion formulae between CFU/FW and CFU/mL_xyl_:

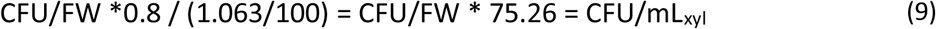

Finally, we can straightforwardly estimate a conversion between CFU/FW and OD or biomass concentration in mg/L_xyl_ using the formulae:

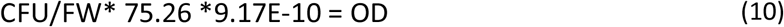

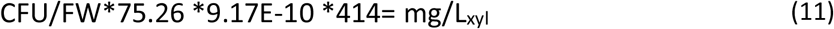

10^9^CFU/gFW thus represents an OD of 69.01, which is equivalent of biomass concentration of 28.57 g/L. 10^10^CFU/gFW thus represents an OD of 690.1, which is equivalent of biomass concentration of 285.7 g/L.

## Acknowledgement

Léo Gerlin was funded by a PhD grant from the French Ministry of National Education and Research. Antoine Escourrou was funded by the French Laboratory of Excellence TULIP (ANR-10-LABX-41 and ANR-11-IDEX-0002-02). The study was funded by the French Laboratory of Excellence (LABEX) project TULIP (ANR-10-LABX-41 and ANR11-IDEX-0002-02). The funders had no role in study design, data collection and analysis, decision to publish, or preparation of the manuscript. We acknowledge the MetaToul platform (Metabolomics & Fluxomics Facilities, Toulouse, France, www.metatoul.fr), which is part of the MetaboHUB- ANR- 11- INBS- 0010 national infrastructure (www.metabohub.fr) and its staff: Cécilia Berges, Edern Cahoreau, and Lindsay Peyriga for access to NMR facilities.

## Competing interest statement

The authors declare no competing interests.

## Authors contributions

Caroline Baroukh: Conceptualization (lead); Data curation (lead); Formal analysis (lead); Funding acquisition (lead); Investigation (equal); Methodology (lead); Project administration (lead); Resources (equal); Supervision (lead); Writing – original draft (lead); Writing – review and editing (equal). Leo Gerlin: Investigation (equal); Writing – review and editing (supporting). Antoine Escourrou: Investigation (equal); Stéphane Genin: Conceptualization (supporting); Methodology (supporting); Resources (equal); Writing – review and editing (supporting).

## Data and code availability

All data and scripts used in this study are available in a Github repository: https://github.com/cbaroukh/MacroscopicModel_XMM_and_InPlanta [public upon publication acceptance]

## Supporting Information

**Figure S1:**
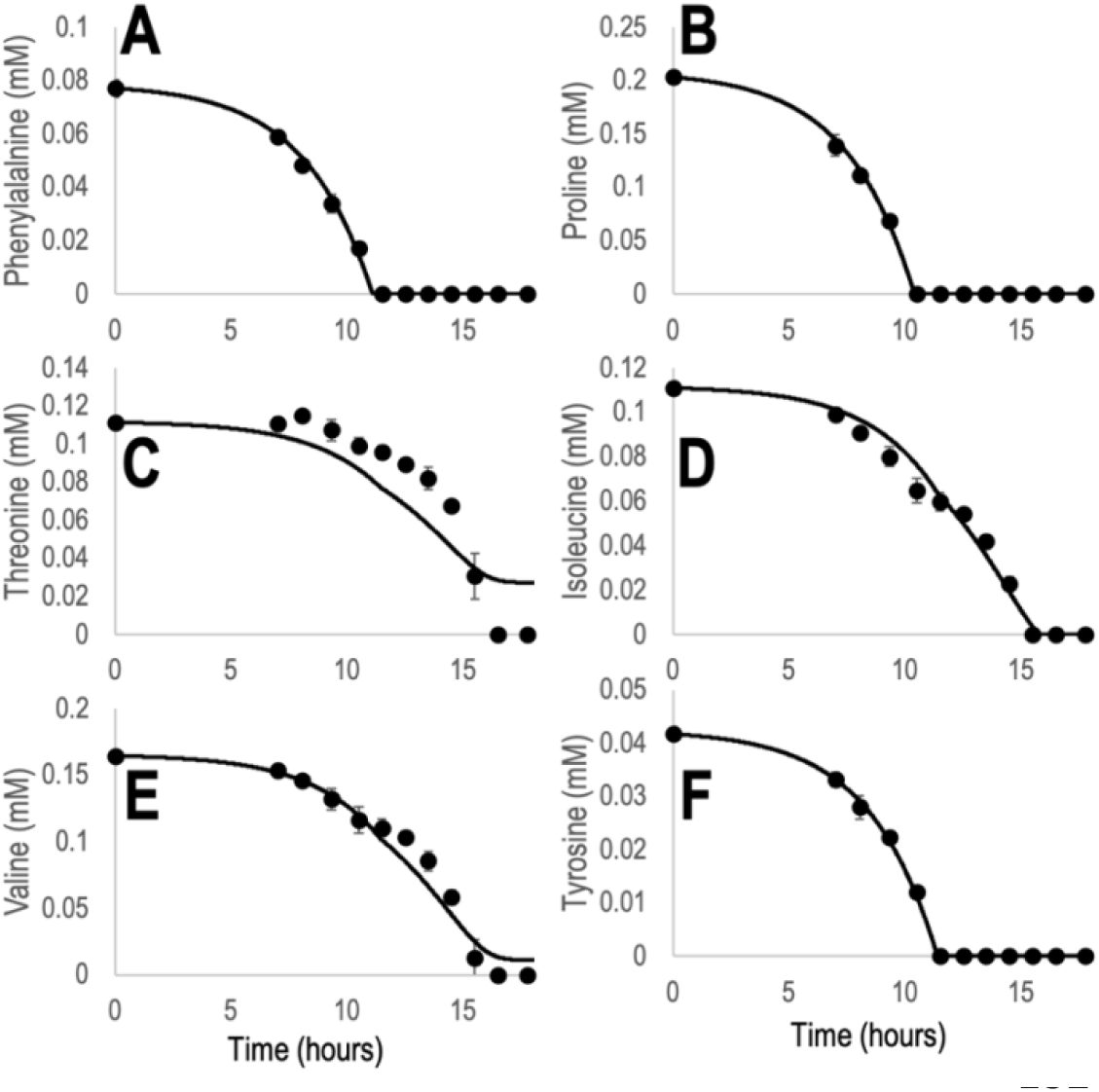
Growth monitoring of *R. pseudosolanacearum* GMI1000 strain in a xylem mimicking media. Dots are experimental data, line represents model’s simulation results. Metabolite concentration over time was assessed by NMR analysis of culture filtrate. A. Phenylalanine. B. Proline. C. Threonine. D. Isoleucine. E. Valine. F. Tyrosine.

**Fig S2:**
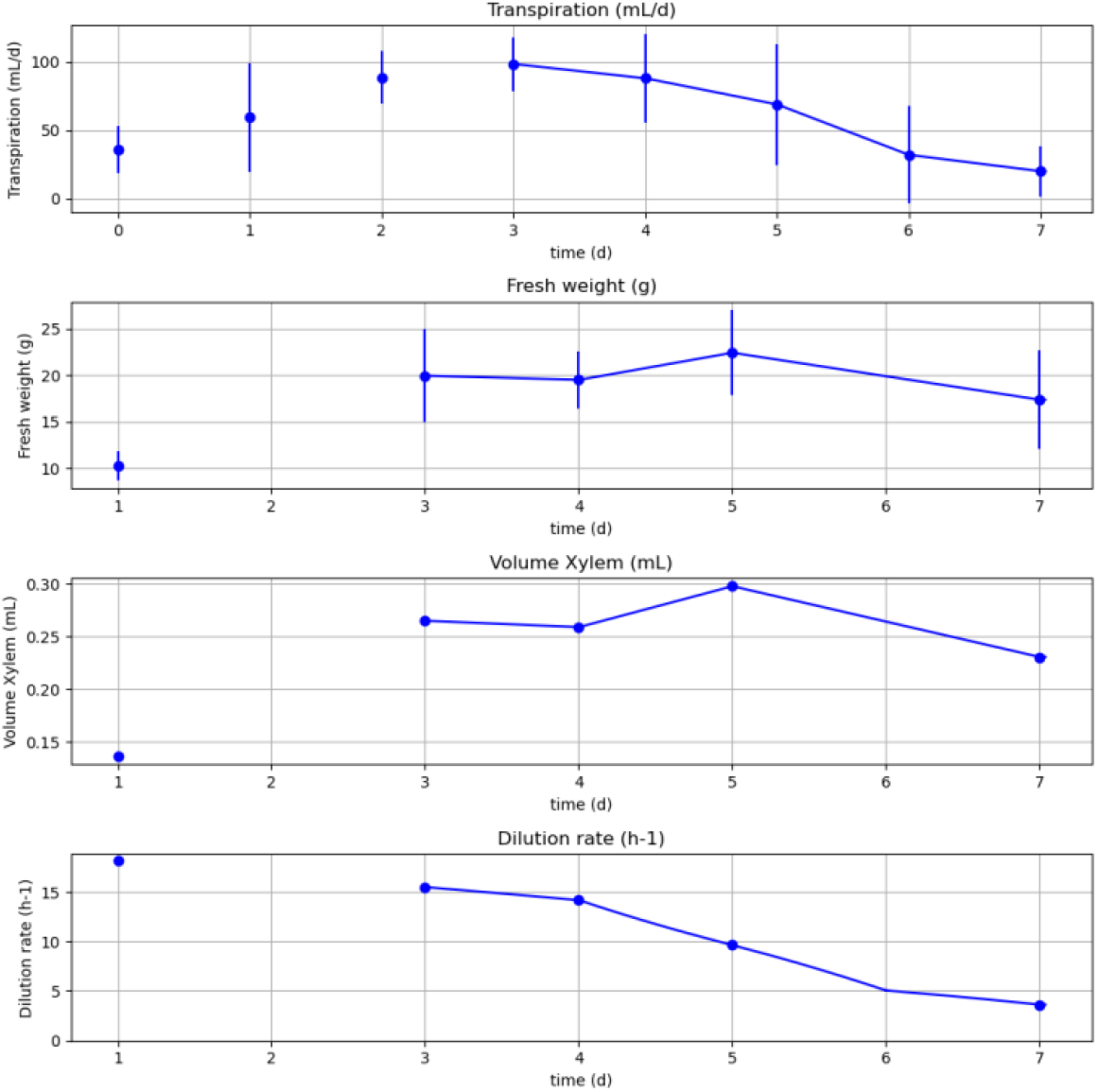
Transpiration, plant fresh weight, xylem volume and dilution rate computed over time. Dots represent values experimentally measured or computed from experimental data. Lines represent interpolated data for models’ simulations.

## Notes

https://github.com/cbaroukh/MacroscopicModel_XMM_and_InPlanta

### Competing Interest Statement

The authors have declared no competing interest.

